# Cortical connectivity supports motoric synchronization to both auditory and visual rhythms in a frontal-temporal network

**DOI:** 10.1101/2024.09.17.613394

**Authors:** Yuhan Lu, Yanlin Yu, Xiaosha Wang, Lang Qin, Jia-Hong Gao, Yanchao Bi, Xing Tian, Nai Ding

## Abstract

Synchronizing motoric responses to metrical sensory rhythms is key to social activities, e.g., group singing and dancing. It remains elusive, however, whether there is a common neural network for motoric synchronization to metrical rhythms from different sensory modalities. Here, we separate sensorimotor responses from basic sensory responses by combining a metrical sensorimotor synchronization task with frequency-domain magnetoencephalography (MEG) analysis. A common frontal-temporal network, not including visual cortex, is observed during both visual- and auditory-motor synchronization, and the network remains in congenitally deaf participants during visual-motor synchronization, suggesting the network is formed by intrinsic cortical connections instead of auditory experience. Furthermore, activation of the left and right frontal-temporal areas, as well as the ipsilateral white matter connection, separately predict the precision of auditory and visual synchronization. These results reveal a common but lateralized frontal-temporal network for visual- and auditory-motor synchronization, which is generated based on intrinsic cortical connections.

## Introduction

Synchronizing motor activity to external sensory stimuli is fundamental to the survival of animals and their social interactions [1–5]. For example, when hearing a loud sound, humans and animals immediately turn their heads towards the sound source [6, 7]. During social interactions, e.g., for males to attract females, a group of frogs can synchronize their choruses based on auditory feedback and fiddler crabs can synchronize their claw movement based on visual feedback [8]. A particular type of sensorimotor synchronization, i.e., synchronizing movement to a rhythmic sensory stimulus, is especially critical for social bonding and communications for humans [8–13]. For example, humans can sing or dance to a beat (i.e., synchronizing vocal or body movements to an auditory rhythm) and sing or play instruments according to the visual movements of a conductor (i.e., synchronizing vocal or body movements to a visual rhythm). Furthermore, a ubiquitous phenomenon during rhythm perception is that one can distinguish strong and weak beats and group beats into meters [14, 15]. The meter-level rhythm can be conveyed by acoustic cues, e.g., sound intensity, but can also be internally tracked without acoustic cues [16–18]. In other words, even when the same auditory and visual item repeats at a constant rate, one can imagine or tap to a 1:2 (march), 1:3 (waltz), or 1:4 meter [19].

Although humans can synchronize movements to sensory rhythms from different modalities [20], they can better synchronize to auditory rhythms [21, 22] – It is well-established that human participants can tap more precisely and more predictively to an auditory rhythm, compared with a visual rhythm [19, 23, 24], a phenomenon not observed in other species [25]. To explain the asymmetry across auditory and visual modalities, it has been hypothesized that since the anatomical connection between auditory and motor areas is stronger than the connection between visual and motor areas [26, 27], visual rhythms are transformed into auditory representations to synchronize movements [28, 29]. This hypothesis is referred to as the auditory-imagery hypothesis. Consistent with this hypothesis, a frontal-temporal network is activated during both visual- and auditory-motor synchronization, including the left inferior frontal gyrus (IFG), bilateral superior temporal gyrus (STG), and left-hemisphere neural fibers connecting IFG and STG [30–32].

Other studies, however, suggest that it is not auditory processing per se that mediates motor synchronization to a visual rhythm. For example, congenitally deaf individuals without any auditory experience can still synchronize to a visual rhythm, and their performance can even be better than hearing individuals [33]. Furthermore, previous studies hint that the neural networks for auditory and visual-motor synchronization separately lateralized to the left [34–36] and right [37–39] hemispheres, which cannot be easily explained if auditory imagery is the only factor driving the sensorimotor synchronization network. Based on these findings, we hypothesize that the neural network engaged by visual- or auditory-motor synchronization task is determined by its intrinsic neural connections with sensory and motor areas in the brain instead of auditory experience (see ref. [40, 41] for similar discussions in the domain visual object recognition). According to this neural-connection hypothesis, auditory imagery may be a consequence or by product if the sensorimotor network overlaps with the neural networks generating auditory imageries.

The current study aims to map the cortical networks for visual- and auditory-motor synchronization. According to the auditory-imagery hypothesis, there is a common network for visual- and auditory-motor synchronization. According to the neural-connection hypothesis, the networks may differ across modalities but should remain in congenitally deaf participants who cannot use auditory experience to mediate visual-motor synchronization. Mapping potentially modality-specific networks for sensorimotor synchronization, however, is challenging – For example, if a visual area is activated during visual- but not auditory-motor synchronization, it is not clear whether the area encodes the visual input or is engaged in visual-motor synchronization. Due to this challenge, most studies focus on modality-general areas for sensorimotor synchronization [42–46]. Here, to overcome the challenge, we design a metrical synchronization task that can separate the neural responses related to sensory and sensorimotor processing in the frequency domain. In the task, participants subvocally count a sequence of auditory or visual items. Critically, the participants counted in cycles of four (**Fig. 1**), creating a sensorimotor rhythm is 1/4 of the sensory rhythm. Therefore, neural responses engaged in basic sensory processing and sensorimotor synchronization are tagged with different rhythms. Additionally, we adjust the speed of rhythm to match the difficulty of auditory and visual synchronization tasks so that a modality-specific effect cannot be explained by task difficulty.

**Figure 1.**
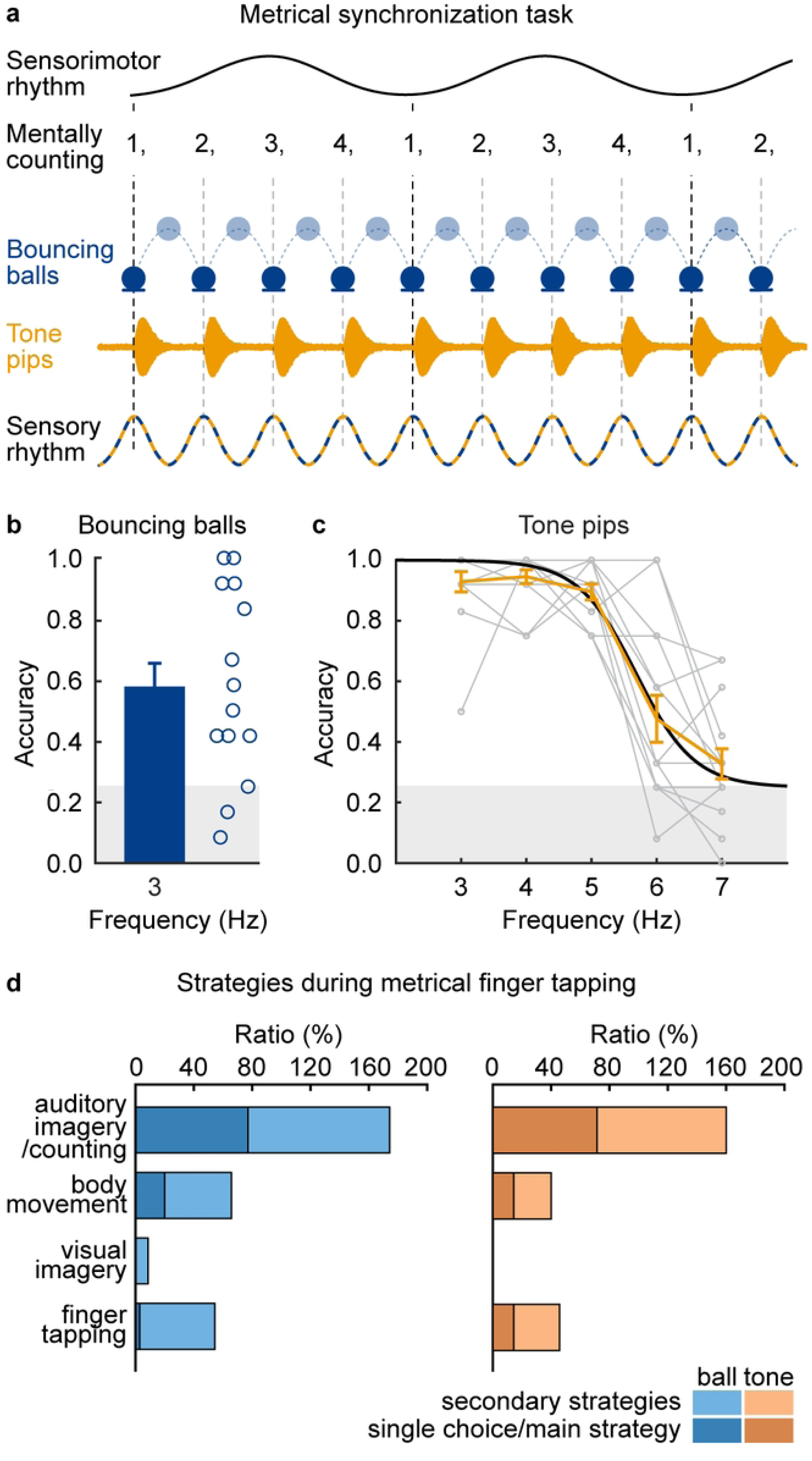
Metrical tracking of auditory and visual rhythms. **a**, Behavioral experiment 1. The experiment presents isochronous visual or auditory sequences and the participants are asked to subvocally count the number of tones or ball bouncing in a cyclic manner (i.e., 1, 2, 3, 4, 1, 2, …) and report the last count. The visual experiment presents a silent video showing a ball dropping and bouncing at a rate of 3 Hz. The auditory experiment presents tone-pips at different rates between 3 and 7 Hz. **b**, Ratio of trials correctly reporting the last count in the visual experiment (*N* = 14). Each circle represents a participant. **c**, Ratio of trials correctly reporting the last count in the auditory experiment as a function of stimulus rates (*N* = 15). In **b** and **c**, participant-averaged accuracy is shown in orange, fitted sigmoid function is shown in black, and each gray dot represent a participant. Error bars represent 1 SEM across participants. **d**, Behavioral experiment 2. The experiment presents the same auditory and visual sequences but participants are asked to tap once every four visual or auditory items (*N* = 35). Survey about the strategies shows that auditory imagery/counting is reported as the most commonly-used strategy.

In the experiments, we recorded neural the neural response to visual or auditory rhythms using MEG, while the participants performed the metrical subvocal counting task. We extracted the neural responses from bilateral IFG, bilateral STG, and visual cortex using source-space imaging and analyzed which cortical areas tracked the sensory or sensorimotor rhythms. Furthermore, we investigated whether the same cortical areas were engaged during visual-motor synchronization in congenitally deaf individuals. We also utilized diffusion tensor imaging (DTI) to probe the which white-matter fiber tracts relate to sensorimotor neural activity or the behavioral performance.

## Results

### Subvocal tracking of auditory and visual rhythms

We developed a subvocal metrical counting task in which the rhythm of sensorimotor activity was dissociated with the rhythm of the sensory input. Specifically, the experiment presented an isochronous sequence of auditory or visual items, and the participants were asked to mentally count each item in a cyclic way, i.e., “1, 2, 3, 4, 1, 2, 3, 4, 1, 2…” (**Fig. 1a**). In this task, imagined vocal activity synchronized to every group of four sensory items, which can be viewed as a meter, and therefore the rhythm of sensorimotor synchronization that is 1/4 of the rhythm of the sensory input. To quantify the accuracy of the subvocal metrical counting task, the participants were asked to report their last count, i.e., a number between 1 and 4, at the end of a sequence. To avoid a response bias, we adjusted the number of items in the sequences so that the last count was uniformly distributed between 1, 2, 3, and 4. Correctly answering the question required the participants to successfully synchronize their counting to every single sensory item throughout the whole sequence.

The visual-motor synchronization task presented a video in which a ball dropping and bouncing 3 times a second while the participants performed the subvocal metrical counting task (**Fig. 1a**). Behavioral accuracy of task was 58% ± 8%, significantly higher than chance (*p* = 0.017, paired two-sided bootstrap, FDR corrected; **Fig. 1b**). In the auditory-motor synchronization task, we presented an isochronous sequence of tone pips and adjusted the speed of the sequence to match the difficulty of the auditory-motor and visual-motor synchronization tasks. Accuracy of the auditory-motor synchronization task decreased with a faster rhythm (**Fig. 1c**). When tone pips were presented at 6 Hz, the behavioral accuracy was 54% ± 7%, roughly matching the accuracy of the visual synchronization task. Therefore, in the following MEG experiment, to match the task difficulty across auditory and visual modalities, the auditory rhythm was presented at 6 Hz and the visual rhythm was presented at 3 Hz. An additional condition, i.e., a 3-Hz auditory rhythm condition, was also included to match the stimulus rate across auditory and visual modalities. When the stimulus rhythm was at 3 or 6 Hz, sensorimotor synchronization occurred at 0.75 or 1.5 Hz, which was within the optimal frequency range for sensorimotor synchronization.

Since the metrical subvocal counting task is not commonly used to study sensorimotor synchronization, we ran an additional experiment to investigate whether it could reflect the core mechanism in well-established sensorimotor tasks, i.e., metrical tapping. The experiment recruited a separate group of participants who were instructed to tap once every four visual or auditory items. A survey after the experiment suggested that most participants spontaneously used strategies similar to subvocal counting even in the metrical tapping task (**Fig. 1d**). Therefore, the metrical subvocal counting task could potentially reveal common mechanisms during metrical sensorimotor synchronization.

### Cortical tracking of sensory and sensorimotor rhythms

The MEG experiment included three types of stimuli, i.e., a 3-Hz visual sequence, a 3- Hz auditory sequence, and a 6-Hz auditory sequence (**Fig. 2a**), and the participants performed the same metrical synchronization task in the MEG experiment. It was confirmed that the task accuracy during the MEG experiment was comparable for the 3-Hz visual and 6-Hz auditory conditions (*N* = 23; *p* = 0.774; Friedman’s chi-square test, FDR corrected). In both conditions, however, there was great variability across participants, with individual task accuracy varying from below 20% to above 80% (**Fig. 1b** and **1c**). For the 3-Hz auditory condition, the mean accuracy was much higher (*p* = 0.00001 and 0.0002, compared with the 3-Hz visual and 6-Hz auditory conditions, respectively; Friedman’s chi-square test, FDR corrected).

**Figure 2.**
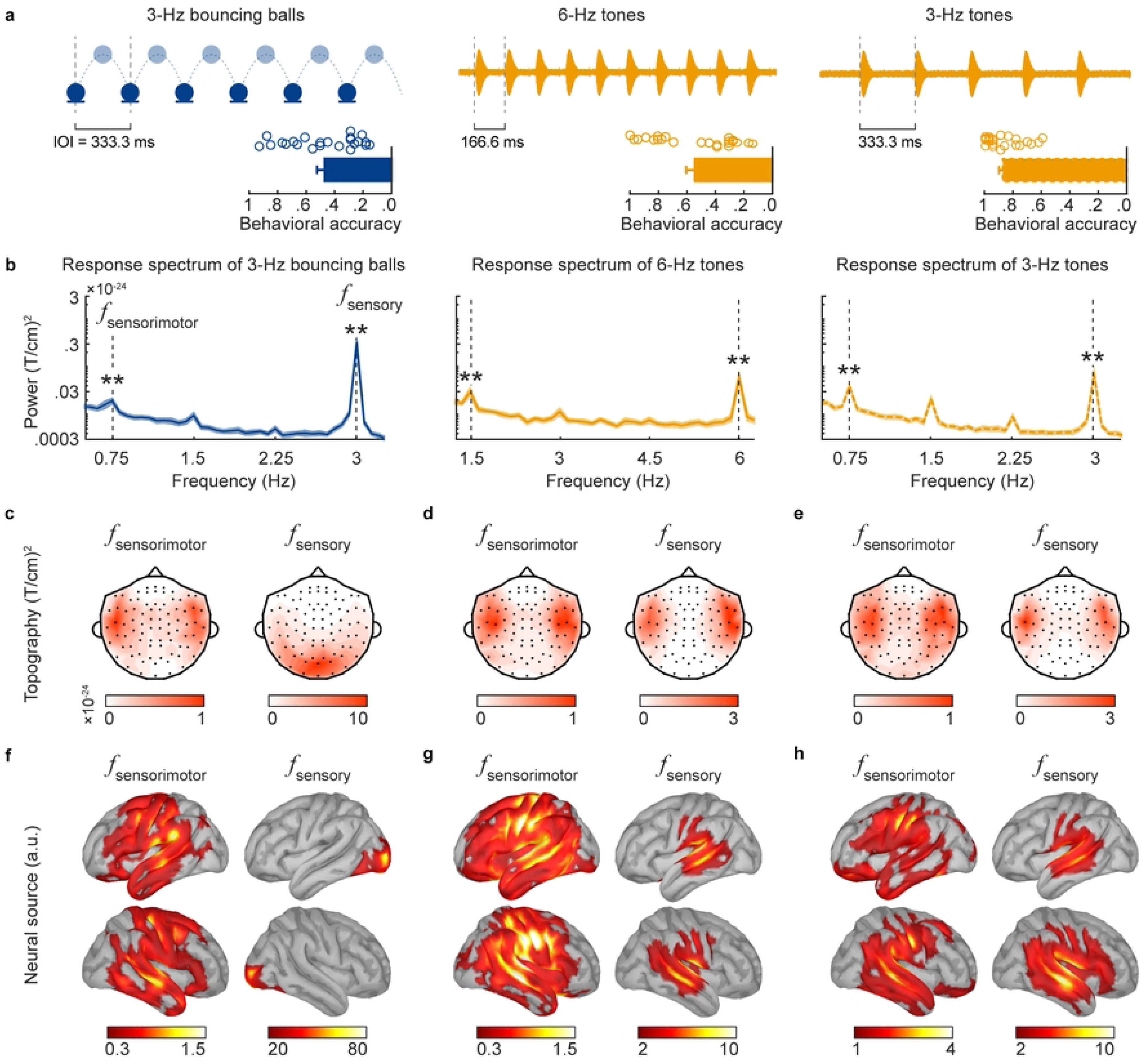
Cortical tracking of sensory and sensorimotor rhythms. **a**, MEG experiment presents three conditions in three separate blocks. In each condition, the mean behavioral accuracy is shown by the bar and each dot denotes one participant. Error bars represent 1 SEM across participants. **b,** The MEG spectrum shows significant peaks at the sensory and sensorimotor rates. The shaded area covers 1 SEM across participants (*N* = 23) on each side, evaluated by bootstrap. \*\**p* < 0.005 (one-sided bootstrap, FDR corrected) **c-e**, Topography of the MEG response at the sensory and sensorimotor rates. Only data from gradiometers (positions shown by black dots) are shown. **f-h**, Source-space dSPM value of MEG responses.

Taking advantage of the metrical synchronization task, we dissociated sensory-rate neural activity, i.e., neural activity tracking individual auditory/visual items presented at 3 or 6 Hz, and sensorimotor-rate neural activity, i.e., neural activity tracking the 3/4 Hz or 6/4 Hz subvocal counting. In the channel-averaged MEG response spectrum, we observed peaks at both the sensory and the sensorimotor rates in all three conditions (*N* = 23; *p* = 0.0001 for all sensorimotor and sensory responses, paired one-sided bootstrap, FDR corrected; **Fig. 2b**). At the sensory rate, the channel-averaged power was significantly stronger in the visual than auditory conditions (*p* = 0.0003 for both 3-Hz and 6-Hz auditory conditions, respectively, paired two-sided bootstrap, FDR corrected). At the sensorimotor rate, the channel-averaged power in the 3-Hz auditory condition, which had higher behavioral accuracy, was significantly stronger than the channel-averaged power in the 3-Hz visual and 6-Hz auditory conditions (*p* = 0.0006 and 0.048, respectively, paired two-sided bootstrap, FDR corrected).

For the visual condition, the sensory-rate response was source localized to the visual cortex in occipital lobe (**Fig. 2c** and **2f**, right panels). For the auditory conditions, the sensory-rate response was localized to areas around bilateral auditory cortex (**Fig. 2d-e** and **2g-h**, right panels). In sharp contrast to the modality-specific sensory-rate responses, the sensorimotor-rate response was source localized to a bilateral temporo-frontal network in all three conditions (**Fig. 2c-h**, left panels), suggesting a modality-general network for sensorimotor transformation that is separated from basic sensory cortices.

### Neural correlation of auditory and visual synchronization accuracy

Next, we investigated the relationship between individual behavior and neural activity, i.e., whether individual behavioral performance could be explained by neural activation in five regions of interest (ROIs), i.e., left IFG, right IFG, left STG, right STG, and visual cortex. We chose the first four ROIs since they showed strong activation in the source space analysis (**Fig. 2f-h**) and the visual cortex since it was also an obvious candidate for visual rhythm processing. The neural response spectrum was averaged within each ROI. For the first four ROIs, the peaks at the sensorimotor rate (shown in **Fig. 3a-d**, lower panels) and the sensory rate were statistically significant (*p* <0.003 for all conditions, paired one-sided bootstrap, FDR corrected). The visual cortex ROI, however, did not show a significant sensorimotor-rate response (*p* > 0.078 for all conditions; **Fig. 3e**).

**Figure 3.**
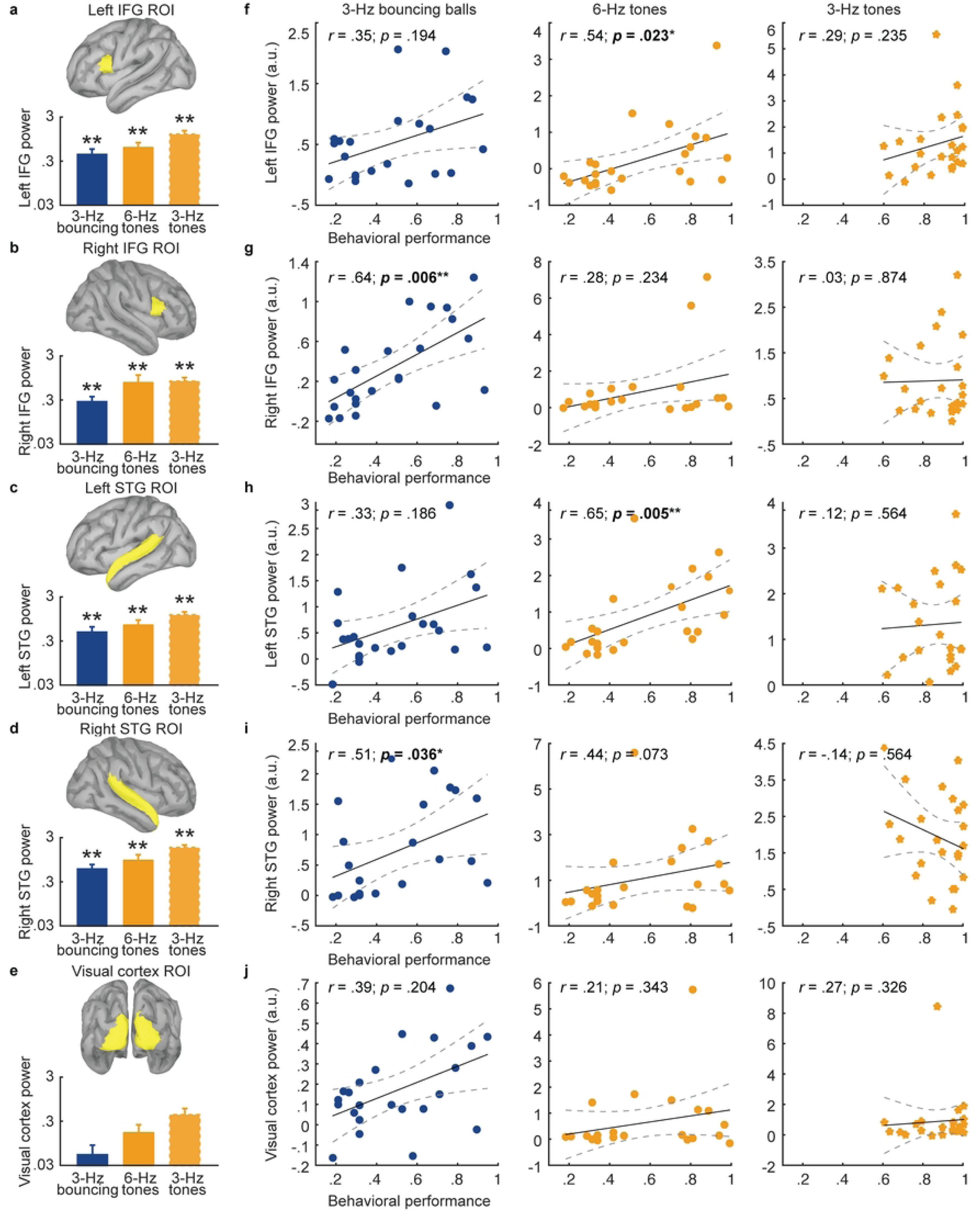
Sensorimotor-rate response power in IFG and STG correlates with individual behavioral performance. a-e, Sensorimotor-rate response power averaged across dipoles within each ROI. Significant sensorimotor-rate response is observed in the first four ROIs, i.e., panels **a-d**. Error bars represent 1 SEM across participants. **f-j,** Relationship between the sensorimotor-rate response power in each ROI and behavioral accuracy in each stimulus condition. The two left-hemisphere ROIs correlate with the performance in the auditory task, while the two right-hemisphere ROIs correlate with the performance in the visual task. Each dot represents a participant and the color indicates stimulus modality. Dashed lines denote 95% confidence interval. \**p* < 0.05, \*\**p* < 0.005.

The sensory-rate power in none of the ROIs correlated with the behavioral performance in either condition (*p* > 0.539, Spearman correlation, FDR corrected). The relationship between sensorimotor-rate power in each ROI and behavioral performance was shown in **Fig. 3f-j**. The left IFG and left STG power correlated with the behavioral performance in the 6-Hz auditory condition (*N* = 23; *p* = 0.023 and 0.006 for IFG and STG, respectively, Spearman correlation, FDR corrected), while the right IFG and right STG power correlated with the behavioral performance in the 3-Hz visual condition (*p* = 0.005 and 0.036, Spearman correlation, FDR corrected).

To further quantify the lateralization effect, we calculated a lateralization index for both IFG and STG, which was the correlation coefficient in the left hemisphere subtracting the correlation coefficient in the right hemisphere, for each modality. The lateralization index in IFG was significantly higher than 0, i.e., showing left lateralization, in the 6-Hz auditory condition and was significantly lower than 0, i.e., showing right lateralization, in the 3-Hz visual condition (*p* = 0.030 for both auditory and visual conditions, one-sided bootstrap, FDR corrected). No significant lateralization was observed for the STG in any condition. These results suggested that the precision of motoric synchronization to auditory and visual rhythms was separately controlled by left- and right-lateralized IFG activation.

### White matter pathways for auditory and visual synchronization

Previous studies have shown that the microstructure of white matters is associated with sensorimotor synchronization [34, 36]. To further probe the potential structural basis of lateralized frontal-temporal networks for auditory and visual synchronization, we acquired diffusion tensor imaging (DTI) data from the same cohort of participants in the MEG experiment (*N* = 23). Specifically, we asked whether the arcuate fasciculi, connecting the inferior frontal and superior temporal areas, contributed to the lateralized cortical responses and behavioral performance. The behavioral accuracy of individual participants roughly fell into a bimodal distribution (**Fig. 4a**), and was fitted by a two-cluster Gaussian model. Based on the fitted clusters, we separated the participants into high performers (*N*_high_ = 7) and low performers (*N*_low_ = 11) and compared the connectivity strength of the arcuate fasciculus between the two groups. The strength of arcuate fasciculus significantly differed between the two groups, with higher fractional anisotropy (FA) value for the low performers than the high performers (*p* = 0.0008 and 0.001 for the right and left arcuate fasciculus, respectively, paired two-sided bootstrap, FDR corrected; **Fig. 4b** and **d**). Importantly, the FA of arcuate fasciculus significantly correlated with the sensorimotor-rate power in the auditory/visual synchronization task: The sensorimotor-rate power of right IFG correlated with the FA of right arcuate fasciculus in the 3-Hz visual condition (*r* = −0.64, *p* = 0.011, Spearman correlation, FDR corrected; **Fig. 4c)**, but not in the auditory conditions. In contrast, the left IFG correlated with the FA of left arcuate fasciculus in the 6-Hz auditory condition (*r* = −0.53, *p* = 0.049, Spearman correlation, FDR corrected; **Fig. 4e**), but not in the other conditions. These results suggested that left/right IFG activation during the auditory/visual synchronization task was constrained by the left/right arcuate fasciculi.

**Figure 4.**
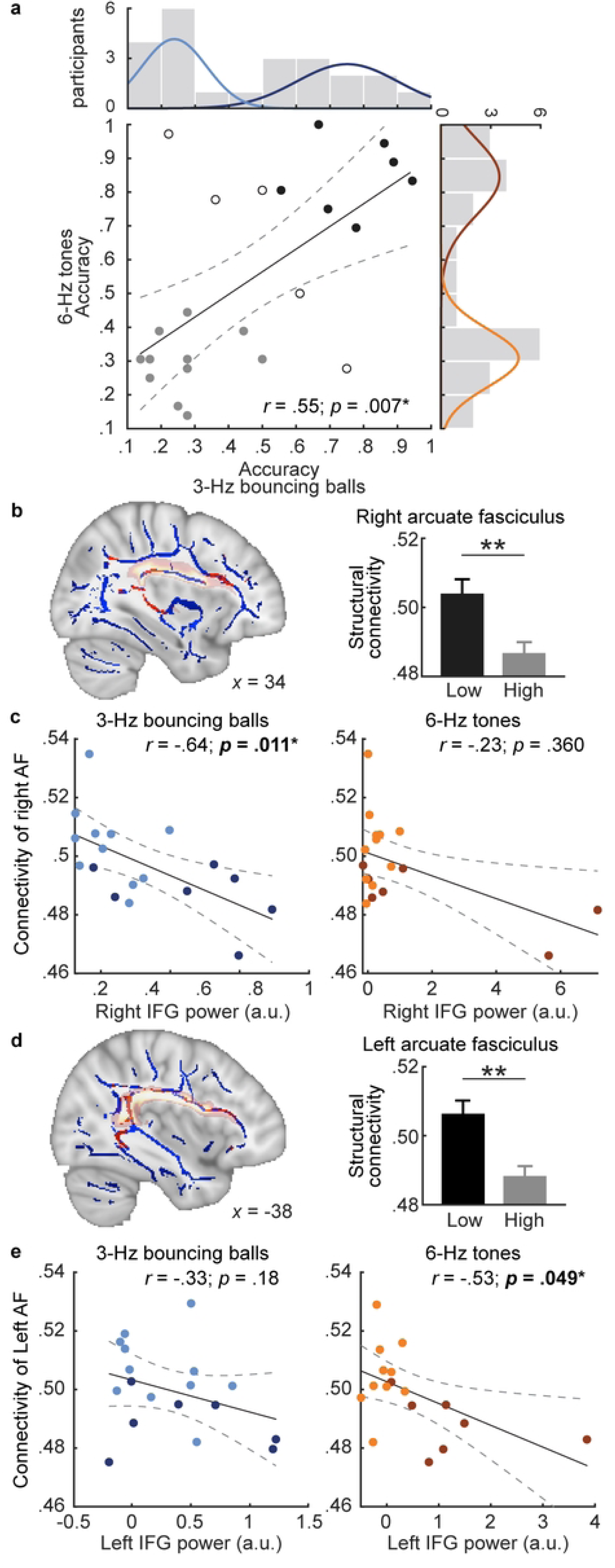
Structural correlates to behaviors and sensorimotor activities. **a**, Histograms of behavioral performance (*N* = 23) are fitted using a two-cluster Gaussian model and the two clusters are separately shown by the darker and lighter curves. Participants are separated based on the two clusters into low and high performers (shown by gray and black circles respectively), who are analyzed in subsequent DTI analyses. **b** and **d**, Sagittal maps of FA generated using tract-based spatial statistics (TBSS). White matter clusters significantly differentiating high and low performers (*N*_high_ = 7, *N*_low_ = 11; *p* < 0.001, two-sided bootstrap, FDR corrected) are shown in red. Areas shaded in light red indicate the template of arcuate fasciculi. **c** and **e**, Scatterplots (*N* = 18) showing the correlation of FA value with sensorimotor-related power over IFG. Each dot is one participant (darker and lighter colors for high and low performers). Error bars represent 1 SEM across participants. Dashed lines denote 95% confidence interval. \**p* < 0.05, \*\**p* < 0.005.

### Visual-motor synchronization in congenital deaf individuals

Previous experiments established that visual-motor and auditory-motor synchronization engaged a common temporal-frontal network, but it remains unclear whether the common network is driven by auditory imageries, i.e., the auditory-imagery-based hypothesis, or anatomical connections to sensory and motor areas, i.e., the neural-connection-based hypothesis. To distinguish these two hypotheses, we conducted the same visual MEG experiment on congenitally deaf participants (*N* = 5) who lacked auditory experience, which was the basis of auditory imagery. The experiment presented the same 3-Hz visual sequence used in the previous experiment and asked the congenitally deaf participants to imagine sign gestures in cycles of fours (**Fig. 5a**). The MEG spectrum showed a significant peak at sensory rate (*p* = 0.0001, paired one-sided bootstrap, FDR corrected; **Fig. 5b**), which was clustered over visual cortex (**Fig. 5c**, lower panels). Critically, a significant peak was also detected at sensorimotor rate (*p* = 0.0008, paired one-sided bootstrap, FDR corrected), which was localized to bilateral frontal-temporal areas (upper panels). These results suggested that the frontal-temporal network for sensorimotor synchronization was independent of auditory processing and therefore strongly supported the neural-connection-based hypothesis.

**Figure 5.**
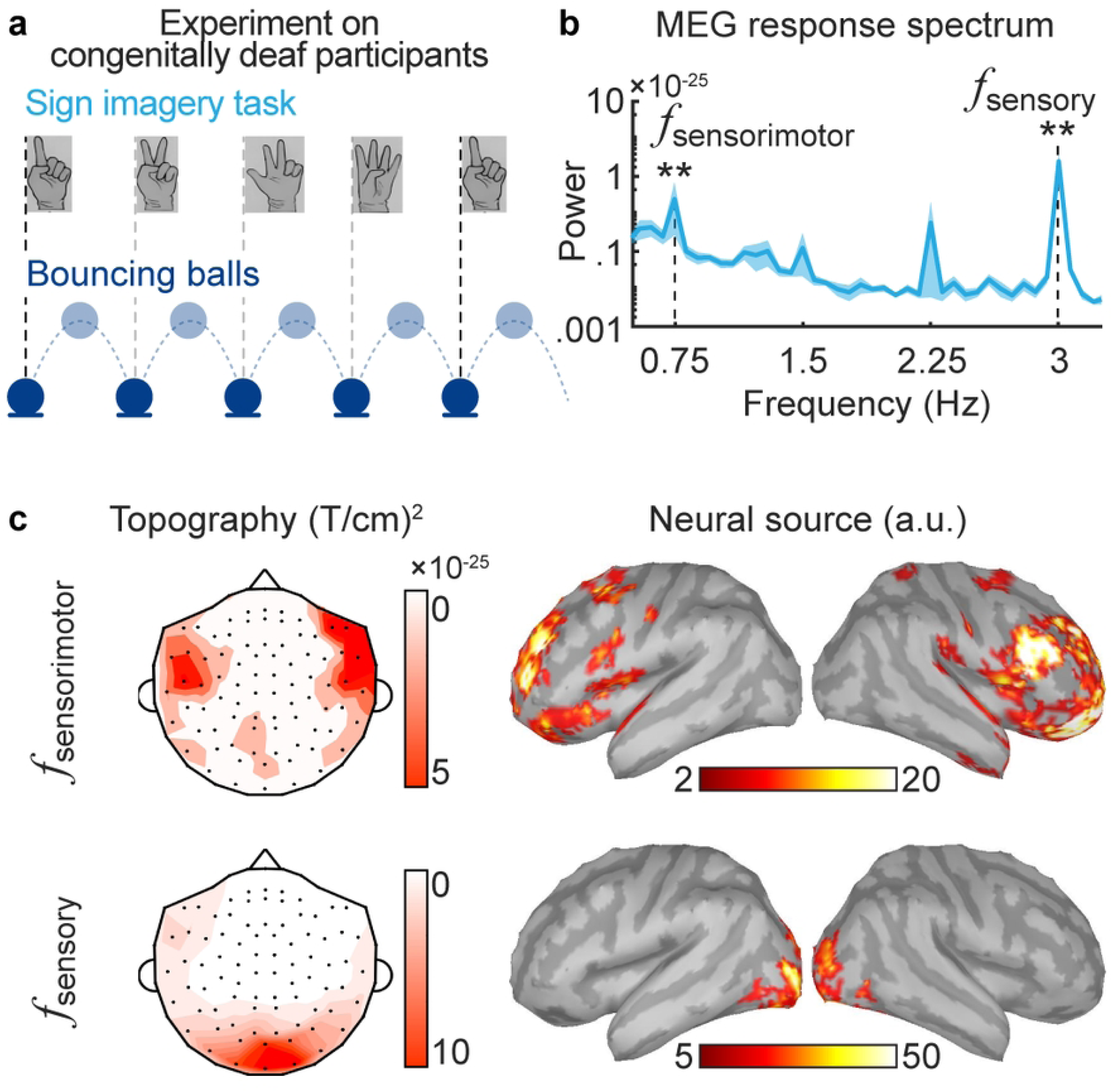
Cortical response during visual-motor synchronization for congenitally deaf individuals. **a**, Congenitally deaf participants (*N* = 5) are asked to count the number of ball bouncing in a cyclic way by imagining sign gestures, i.e., 1, 2, 3, 4, 1, 2,… **b**, The MEG spectrum shows significant peaks at both the sensory and sensorimotor rates. The shaded area covers 1 SEM across participants on each side, evaluated by bootstrap. **c**, Topography and source map of the MEG response at the sensory and sensorimotor rates. Left: gradiometers (black dots) are shown. Right: dSPM values are shown. \*\**p* < 0.005.

## Discussion

The current study demonstrates that motoric synchronization to auditory and visual rhythms both engage a frontal-temporal network, although the network is separately lateralized to the left and right hemispheres for the auditory and visual inputs. Critically, results from congenitally deaf participants demonstrate that the frontal-temporal network for visual-motor synchronization is determined by intrinsic neural connections instead of being mediated by auditory processing.

Here, we distinguish neural activity related to sensorimotor processing from neural activity encoding the basic sensory input in the frequency domain, using a metrical synchronization task paired with frequency-domain analysis of MEG. This paradigm does not assume that the same network underlies visual- and auditory-motor synchronization, but the results still reveal similar networks during visual and auditory tasks. Crucially, this paradigm shows that the visual cortex is not engaged in visual-motor synchronization, and is only engaged in encoding the input visual rhythm. The metrical synchronization task relies on the fact that motor activity can flexibly synchronize either to the stimulus rate, e.g., *f*, or rates that are metrically related to the stimulus rate, e.g., *f*/4, *f*/2, and 2*f*… [47, 48]. When the stimulus rhythm is too slow, listeners tend to subdivide the beats and synchronize to a frequency that is a multiple of *f*. When the stimulus rhythm is too fast, listeners can better synchronize to a frequency that is a fraction of *f*. According to previous studies on auditory beat tapping, when the stimulus rate *f* is 3 or 6 Hz, participants can tap well at both *f*/4 and *f* [23]. Similarly, previous studies have also shown that participants can reliably tap to a visual rhythm at 1.67 Hz when the visual rhythm is conveyed by a bouncing ball [33]. Consistent with these finger tapping experiments, our subvocal metrical counting experiment reveals precise sensorimotor synchronization to a 3-Hz visual/auditory sequence and a 6-Hz auditory sequence – The behavioral accuracy in the current experiment does not reflect the accuracy of a single count. Instead, a correct response requires the participants to correctly synchronize to all 44-47 items within a sequence. Therefore, 50% behavioral accuracy indicates no single synchronization error in half of the trials.

Classic studies on metrical sensorimotor synchronization mostly engage a metrical tapping task but according to our survey most participants spontaneously use counting or auditory imagery as a strategy to achieve metrical finger tapping (**Fig. 1d**). In principle, the participants can entrain a slow rhythm at one quarter of the stimulus rate and tap based on entrained slow rhythm. Instead, the survey suggests that the participants imagine hearing or speaking a sound for every sensory item and use the imagined sound to distinguish items to tap to and items that should be skipped, similar to a subdivision strategy. In the survey or even in the subvocal counting task, we do not distinguish imagined speaking and imagined hearing since it is usually difficult to distinguish the two types of imagery. The survey suggests that auditory/speech imagery is a common strategy for metrical sensorimotor synchronization and the findings here can potentially generalize to metrical tapping tasks and other tasks that requires synchronization to a metrical rhythm. Why do the participants tend to spontaneously engage auditory/speech imagery during metrical synchronization? A possibility is that the frontal-temporal network for visual- and auditory-motor synchronization anatomically overlap the neural network for speech imagery [49, 50]. Therefore, the sensorimotor synchronization task may automatically activate the network for auditory/speech imagery or that the participants use auditory/speech imagery to enhance the activation of the sensorimotor network.

Nevertheless, since the frontal-temporal network for visual-motor synchronization is also observed in congenitally deaf participants, it indicates that auditory experience or auditory imagery is not necessary for the activation of the network. Instead, the results suggest that the frontal-temporal network for visual-motor synchronization is caused by the intrinsic white matter connections among sensory and motor areas (i.e., the neural-connection-based hypothesis). The neural-connection-based hypothesis is a strong hypothesis that assumes strong connections between neural anatomy and cognitive function but the hypothesis has received support in other cognitive domains. For example, for sighted individuals, a neural representation of color is found in the anterior temporal lobe, a cortical area that receives input from visual areas. For congenitally blind individuals, however, the same area for color representation is found suggesting that the encoding of color in anterior temporal lobe is not determined by visual experience but is instead determined by intrinsic neural connections, i.e., structure of the brain [41]. Here, we see a very similar case for the network for visual-motor synchronization.

During the metrical tapping task, it is likely that the brain groups every four sensory items into a chunk and synchronize motor activity to the chunks. This kind of chunking, however, relies on a top-down rule, i.e., every four items constructing a chunk, and differs from bottom-up chunking based on stimulus features, i.e., grouping visually similar images into a chunk [51]. Indeed, previous studies have shown, e.g., neural activity in visual cortex can track visual features, as well as the statistical regularity of sequentially presented images [52], but the current study demonstrates that visual cortex only track individual visual items during metrical visual-motor synchronization. The frequency-tagging paradigm here has been previously applied to separate higher-level neural processing, e.g., sentence processing [17, 53, 54], musical meter processing [16], and higher-level patterns in visual sequences [55, 56], from low-level sensory encoding. A recent study has also applied the frequency-tagging paradigm to study the neural basis of auditory imagery, and intracranial EEG recordings demonstrate that auditory-imagery-related activity occurs in the left IFG [57], consistent with the current result that auditory-motor synchronization involves a left-lateralized frontal-temporal network. A limitation of the current MEG-based localization is that MEG is primarily sensitive to cortical sources [58] and therefore future studies may utilize intracranial EEG or more advanced MEG deep-source imaging methods to probe whether subcortical areas, which play critical roles in rhythm processing [45, 59, 60], are engaged in visual- and auditory-motor synchronization.

DTI analysis further demonstrates that the strength of the left and right arcuate fasciculus can predict ipsilateral IFG activation during the auditory and visual synchronization task. Similarly, recent studies that analyze spontaneous overt vocal synchronization to auditory syllables has highlighted that the precision of behavioral auditory-motor synchronization can be predicted by neural activity in the left inferior frontal region and the ipsilateral arcuate fasciculus [34, 61, 62]. Similar to these studies, we observed that the behavioral sensorimotor synchronization accuracy exhibits a bimodal distribution across participants and used this property to demonstrate separable anatomical bases for visual- and auditory-motor synchronization.

In summary, we directly compare neural networks involved in motoric synchronization to auditory and visual rhythms, and control task difficulty across auditory and visual modalities to avoid difficulty-related difference in cortical activation. We observe a frontal-temporal network that is separately lateralized to the right and left hemispheres for visual- and auditory-motor synchronization, and this network is constrained by white-matter connections between temporal and frontal areas, instead of auditory imageries.

## Materials and Methods

### Participants

The auditory and visual behavioral experiments recruited 15 (9 males, 19-28 years old, mean 23 years old) and 14 participants (8 males, 18-26 years old, mean 22 years old), respectively. The finger-tapping behavioral experiment recruited 35 participants (19 males, 19-28 years old, mean 24 years old). The MEG-DTI combined experiment on normal-hearing participants recruited 23 participants (11 males, 18-29 years old, mean 22 years old), and the MEG experiment on congenitally deaf individuals recruited five participants (2 males, 25-31 years old, mean 28 years old). The five deaf participants reported being born profoundly deaf and had deaf parents. The hearing loss (>95 dB SPL) was confirmed by a standard pure-tone audiometry procedure. They used sign language as the primary means of communication. Two deaf participants never used hearing aids, two reported using hearing aids in the past, and one reported occasional use at the time of the experiment. All participants were right-handed [63] and were native Chinese speakers (except deaf participants). Each participate would not attend more than one experiment. The sample size per experiment was predetermined according to previous studies: sample sizes for behavioral and EEG experiments was typically between eight to 22 [16, 23, 64], and the behavioral performance reported here was consistent across experiments (*N* = 92). The experimental procedures were approved by the Research Ethics Committee of the East China Normal University, Zhejiang University, and Peking University, and written informed consent was obtained from each participant before the experiment.

### Auditory stimuli

The stimulus was an isochronous sequence of 1,000-Hz pure tones (**Fig. 1a**). Each sequence consisted of 44 to 47 tones (equally distributed in each experiment). Each tone’s duration was 50 ms, with 10-ms onset and offset cosine ramps. The tones were presented at five different stimulation rates, i.e., 3, 4, 5, 6, and 7 Hz, in the behavioral experiment, and two rates, i.e., 3 and 6 Hz, in the MEG experiment.

### Visual stimuli

The visual stimulus was a ball dropping and bouncing at 3 Hz (**Fig. 1a**). The ball appeared as a white circle (RGB: 170, 170, 170; diameter: 0.3°) in a gray background (RGB: 90, 90, 90). The ball was released from 3.7° above the screen center (i.e., the ‘ground’) and bounced to the same height. The initial velocity of the circle was 0 at the highest point and the acceleration was 0.6°/frame^2^. The size of visual stimulus was chosen so that participants could clearly see the whole visual stimulus when fixating at the center. A bouncing-ball sequence consisted of 44 to 47 repetitions (equally distributed in each experiment), and the refresh rate of the screen was 60 Hz. In addition, in the behavioral experiments, we assessed how the behavioral performance was influenced by different types of visual stimulus, including visual flashes, videos of finger-tapping, rotating-petal, and rotating visual flashes (not reported). The bouncing ball stimulus yielded the best behavioral performance and was used in the MEG experiments.

### Experimental procedures and tasks

#### Behavioral experiments (subvocal counting)

The auditory experiment presented the tone sequence at five different rates, i.e., 3, 4, 5, 6, and 7 Hz, and each rate was presented in a separate block. Each block consisted of 12 sequences and the five blocks were presented in a randomize order. When listening to each sequence, participants were instructed to subvocally count each tone in a cyclic manner (**Fig., 1a**) and report the last count by a button press, i.e., pressing 1, 2, 3, or 4 on a response box using the right hand. After the key press, feedback about correctness was provided and the next trial was presented after a silent interval randomized between 1 and 2 s (uniform distribution). Before the main behavioral experiment, participants received training – Two trials were presented at each of five stimulation rates, and slower rates were presented first. If the participant counted the sequence wrong, they could replay the stimulus. The visual experiment consisted of 12 trials. The task was the same as the auditory experiment, except that participants subvocally counted whenever the ball hit the ground. The participants also received a training section in which three sequences were presented.

#### Behavioral experiment (finger tapping)

The experiment was the same as the first behavioral experiment except that participants were instructed to tap right index finger every four bounces/tones and to report if they tapped to the last one. The instruction carefully avoided words that may trigger other strategies. Participants first performed the visual experiment prior to the auditory experiment to avoid sense of auditory beats during visual experiment [24, 44]. After the visual and auditory experiments, participants completed a survey about the strategies they used during the two experiments. They were provided with four common strategies: tapping with finger, synchronizing with body parts, imagining visual beats, and imagining auditory beats or counting. Participants can choose at least one option and can mark the most suitable one (single choice or main strategy: two marks; secondary strategies: one mark). They could also suggest additional strategies beyond these options. We measured the total marks of each strategy for the experiments (**Fig. 1d**).

#### MEG experiment (normal-hearing individuals)

The MEG experiment presented three types of stimuli, i.e., (1) 3-Hz visual sequence, (2) 3-Hz auditory sequence, and (3) 6-Hz auditory sequence, in three separate blocks. Each block consisted of 36 trials and the three blocks were presented in a randomized order. The participants had a 2-min break after each block. The stimuli and task were the same as the behavioral experiments. Participants were asked to close their eyes in the auditory block, and fixated at a cross at the screen center during the visual blocks. After MEG recording, each participant had T1-weighted and a diffusion-weighted MRI scanning.

Before MEG recording, participants received a training section. The training section consisted of three blocks that were the same as the three blocks in the MEG experiment except that each block consisted of 8 trials and a sequence could be replayed upon request. Participants were informed that during both training and the main experiment, they should not synchronize any part of body with the stimulus or count aloud.

#### MEG experiment (deaf individuals)

The experiment was the same as the MEG experiment on normal-hearing participants except that only the visual condition was presented and the participants counted through sign imageries. After MEG recoding, the participants received T1-weighted MRI imaging.

### MEG data acquisition and pre-processing

Neuromagnetic responses were recorded using a 306-sensor whole-head MEG system (Elekta-Neuromag, Helsinki, Finland) in a magnetically shielded room at Peking University. The MEG signals were sampled at 1 kHz, with a 0.3-Hz high-pass filter applied offline. The system had 102 magnetometer and 204 planar gradiometers. Four head position indicator (HPI) coils were used to measure the head position inside MEG. The positions of three anatomical landmarks (nasion, left, and right pre-auricular points), the four HPI coils, and at least 200 points on the scalp were also digitized before experiment. Temporal Signal Space Separation (tSSS) was used to remove the external interference from MEG signals [65]. To remove ocular and heartbeat-related artifacts in MEG, independent component analysis (ICA) was performed. The response during each trial was extracted, down-sampled to 60-Hz sampling rate.

For MEG source localization, structural T1-weighted data were collected from all participants using a Discovery MR750 3.0T MRI system (GE Healthcare, Milwaukee, WI) at Peking University. A 3-D magnetization prepared rapid gradient echo T1-weighted sequence was used to obtain 1 × 1 × 1 mm3 resolution anatomical images, with following parameters: TR = 2530 ms, TE = 2.98 ms, flip angle = 7°, slice thickness = 1 mm.

### Frequency-domain analysis

In the frequency-domain analysis, to avoid the response to the sound onset, the response during the first two seconds of each trial were removed and the remaining response was 10 s in duration. The average of all trials in the same stimulus condition was transformed into the frequency domain using the Discrete Fourier Transform (DFT) without any additional smoothing window. If the complex-valued DFT coefficient at frequency f was denoted as X(f), the response power was |X(f)|2. The DFT was separately applied to each MEG sensor. Response power from the two collocated gradiometers were always averaged. For channel-averaged analyses, the response power spectrum was averaged across all gradiometers. The response power at the target frequency was further normalized by subtracting the power averaged over four neighboring frequency bins (two bins on each side).

### Source-space analysis

The MEG responses averaged over trials were mapped into source space using cortex constrained minimum norm estimate, implemented in the Brainstorm software [66]. The T1-weighted MRI images were used to extract the brain volume, cortex surface, and innermost skull surface using Freesurfer [67]. Three anatomical landmarks in the MRI images, i.e., nasion, left, and right pre-auricular points, were marked manually. The three anatomical landmarks and digitized head points were used to align the MRI images with MEG sensor array. The forward MEG model was derived based on the overlapping sphere model. The identity matrix was used as noise covariance. Source-space activation was measured by the dynamic statistical parametric map [68]. Individual source-space responses, consisting of 15,002 elementary dipoles over the cortex, was rescaled to the ICBM 152 brain template for source-space analyses. For ROI analysis, a frontal-lobe ROI, i.e., the pars opercularis and its right homolog, was defined according to an automated landmark-based registration algorithm. To illustrating spectrum power, the normalized response power of each dipole was calculated as the power at f minus the power averaged over four neighboring frequency bins around f (two bins on each side). The response power averaged across all dipoles in the ROIs was used to further power and correlation analyses. For illustration, all surface topography were smoothed using SurfStatSmooth at 3 mm.

### Anatomical connectivity analysis

Scanning parameters and diffusion measures. Diffusion-weighted MRI scans were acquired on a 3T MRI scanner (GE Healthcare, MR750, Milwaukee, WI) using a 32-channel head coil, at the Peking University. Diffusion images were acquired with an echo-planar imaging sequence optimized for DTI-MRI of white matter (70 axial slices; TR, 6,000 ms; TE, 64 ms; flip angle, 90°; slice thickness, 2 mm; acquisition matrix, 112 × 112; voxel size, 2 × 2 × 2 mm3, phase-encoding direction: posterior-anterior). 10 unweighted scan and 64 diffusion scans with a b-value of 1,000 s/mm2 were acquired.

### DTI-MRI analysis

Diffusion images was first denoised and removed Gibb’s ringing artefacts in the MRtrix3 software package [69]. Subsequently, eddy current distortions and head motions were corrected by using eddy in FMRIB’s Diffusion Toolbox (FSL) [70]. A quality check was performed, no participant was discarded and the number of outlier slices per participant was below a threshold of 10. Brain extraction was performed by using bet2 command. Diffusion tensors was fitted on eddy-corrected data using dtifit command included in FSL. Finally, FA map for each participant were generated using the eigenvalues extracted from the diffusion tensors, using the JHU DTI-based white matter atlas [71].

### Separation of high and low performers

We used the k-means++ clustering algorithm to separate all participants into two clusters, i.e., the high and low performers, based on the behavioral performance and fitted a Gaussian curve for each cluster.

### Statistical analysis and significance tests

All analyses were conducted using within-participants comparisons and non-parametric methods. We applied FDR correction to all tests. For behavioral performance, we used the Mann-Whitney test for two-condition comparisons and Friedman’s chi-square test for multiple-condition comparisons. For functional and structural brain data, we employed bias-corrected and accelerated bootstrapping [72]. Spearman’s rho was used for correlations between behavioral and brain data.

In the bootstrap procedure, all participants were resampled 10,000 times with replacement. For two-sided comparisons, if the data population in one condition was greater/smaller than A% of the data population in the other condition, the significance level was (200A + 1)/10,001. For one-sided comparisons, if the data population in one condition was greater than A% of the data population in the other condition, the significance level was (100A + 1)/10,001.

#### Spectral peak

The statistical significance of a spectral peak at frequency f was tested by comparing the response power at f with the power averaged over four neighboring frequency bins around f (two bins on each side). In the bootstrap procedure, the MEG waveform was averaged across a group of resampled participants and converted into the frequency domain. The comparison was one-sided. This significance test was only applied to the response at sensory rate and sensorimotor rate.

#### Structural connectivity

A two-sided bootstrap test was used to compare the FA value of white matter fibers between high performers and low performers.

## Acknowledgments

We thank Jiajie Zou, Jianfeng Zhang, Jieyu Lin, Qian Chu, and Lingxi Lu for thoughtful comments on previous versions of the manuscript. Work is supported by National Key Research and Development Program of China 2021ZD0204105, National Natural Science Foundation of China 32222035, and Major Scientific Research Project of Zhejiang Lab 2019KB0AC02 to N.D., and National Natural Science Foundation of China 32071099 and 32271101, Program of Introducing Talents of Discipline to Universities, Base B16018, and NYU Shanghai Boost Fund to X.T.

## Author Contributions

N.D., Y.L., and X.T. designed the experiment. Y.L., X.W., and L.Q. collected the data, Y.L. and Y.Y. performed the analyses, Y.L., N.D., and X.T. drafted, reviewed, and corrected the manuscript. N.D., X.T., Y.B., and J.G. supervised the project.

